# When should a pathogen strike? Navigating the landscape of birth pulses and mating season

**DOI:** 10.64898/2026.10.15.759214

**Authors:** Bhargav D. Srinivasan, Emily L. Bruns, Samuel V. Hulse

**Affiliations:** Department of Biology, University of Maryland, College Park, United States

**Keywords:** host-pathogen, phenology, floral pathogens, infection timing

## Abstract

Susceptibility to pathogens often differs across host life stages, which in turn depends on seasonal host phenology. For many species, juveniles are highly susceptible relative to adults, which given constrained birth periods can lead to seasonal variation in the average host susceptibility. Similarly, for pathogens with complex lifecycles, or seasonally dependent development, peak transmission may be temporally constrained. Maximum pathogen transmission may therefore coincide with a greater abundance of either juvenile or adult hosts. For instance, the birthing season presents a highly susceptible pool of hosts while the mating season presents a higher rate of contact between potentially less susceptible hosts. When these two transmission opportunities are separated in time, it is unclear whether pathogens should evolve a single, intermediate transmission timing or undergo disruptive selection to exploit different host life stages. To investigate how pathogens might navigate this complex landscape, we developed an age-structured model that allows us to test the impacts of host life history on pathogen prevalence and evolution, inspired by a fungal plant pathogen. Parameterizing the model for *Silene latifolia*, we find that disease prevalence is highest when seed germination coincides with flowering, and that separating the two reduces disease. Using invasion analysis, we find that at the estimated *S. latifolia* parameters, a single evolutionarily stable transmission timing closely tracks germination time, with infectious flowering centered about seven weeks after the median germination day. When juveniles mature faster or are much more susceptible than adults, or under parameters for *Dianthus pavonius*, a pathogen genotype that transmits to spring-germinating juveniles and one that transmits later in the year can instead each invade the other and coexist as a polymorphic population when germination is split between spring and summer cohorts. Our model may help explain observed phenological differences seen in wild populations of infected plants.

## Introduction

Many pathogens exhibit strong seasonality, where maximum transmission can be restricted to months or even weeks (Altizer et al. 2006). While seasonal changes in abiotic factors such as temperature and moisture can impact transmission rates, seasonal changes in host demography, behavior, and age-structure can also be important, especially for directly transmitted pathogens (MacDonald & Brisson 2022, Caldwell et al. 2021). Seasonally structured life histories are common, with behaviors such as mating, migration, and hibernation synchronized with environmental cues. How hosts respond to the environment can therefore determine the timing of disease transmission. For example, when host behavior determines pathogen exposure, transmission depends on the same environmental cues (Altizer et al. 2006). Because many of these life-history events recur with seasonal forcing, the transmission opportunities they create are predictable. Pathogens may consequently evolve strategies to alter the timing of host behaviors in ways that enhance transmission (Saadatnia & Golkar 2012, Lovett et al. 2020). These interactions raise a critical evolutionary question: how should pathogens best manipulate host phenology to maximize transmission?

One strategy for pathogens to respond to seasonal variability may be to target a specific host age demographic. Juvenile and adult hosts often differ in susceptibility, contact rates, and transmission pathways (Armitage & Boomsma 2010, Bruns et al. 2017, Duca 1948, Slowinski et al. 2025, Viner et al. 2021, Goldstein et al. 2021) – all of which affect a pathogen’s fitness. Since the relative abundance of juveniles and adults varies across the season, pathogens may face ecological tradeoffs between maximizing transmission at separate host life stages. Two of the most important transmission opportunities are the birthing season and the mating season (Pacala & Dobson 1988, Cornell et al. 2008, Izhar & Ben-Ami 2015). For example, a seasonal spike in susceptible juvenile populations can lead to an increase in transmission, as juveniles typically cannot mount as robust an immune response as adults (van Dijk et al. 2014, Buckingham et al. 2023). Prioritizing juvenile infection might leverage the higher susceptibility of juveniles, but miss out on a larger pool of adult hosts. Conversely, optimizing transmission for adults can increase disease spread via sexual transmission, or greater contact with other hosts and vectors, but is constrained by higher adult resistance. For pathogens with a limited transmission window, the window of peak juvenile abundance may not coincide with the window of peak mating contact, so it is unclear what strategy selection on the pathogen will favor (Figure 1). In some systems, pathogens manipulate host phenology to enhance transmission, for example altering the timing of reproductive events in their hosts (Hafer 2016, Seppälä & Jokela 2008). As such, aspects of host development and phenology, such as maturation rates, susceptibility, or flowering times will determine the temporal segregation of different host-life stages, and the extent to which a pathogen must “choose” which stage to target.

**Figure 1.**
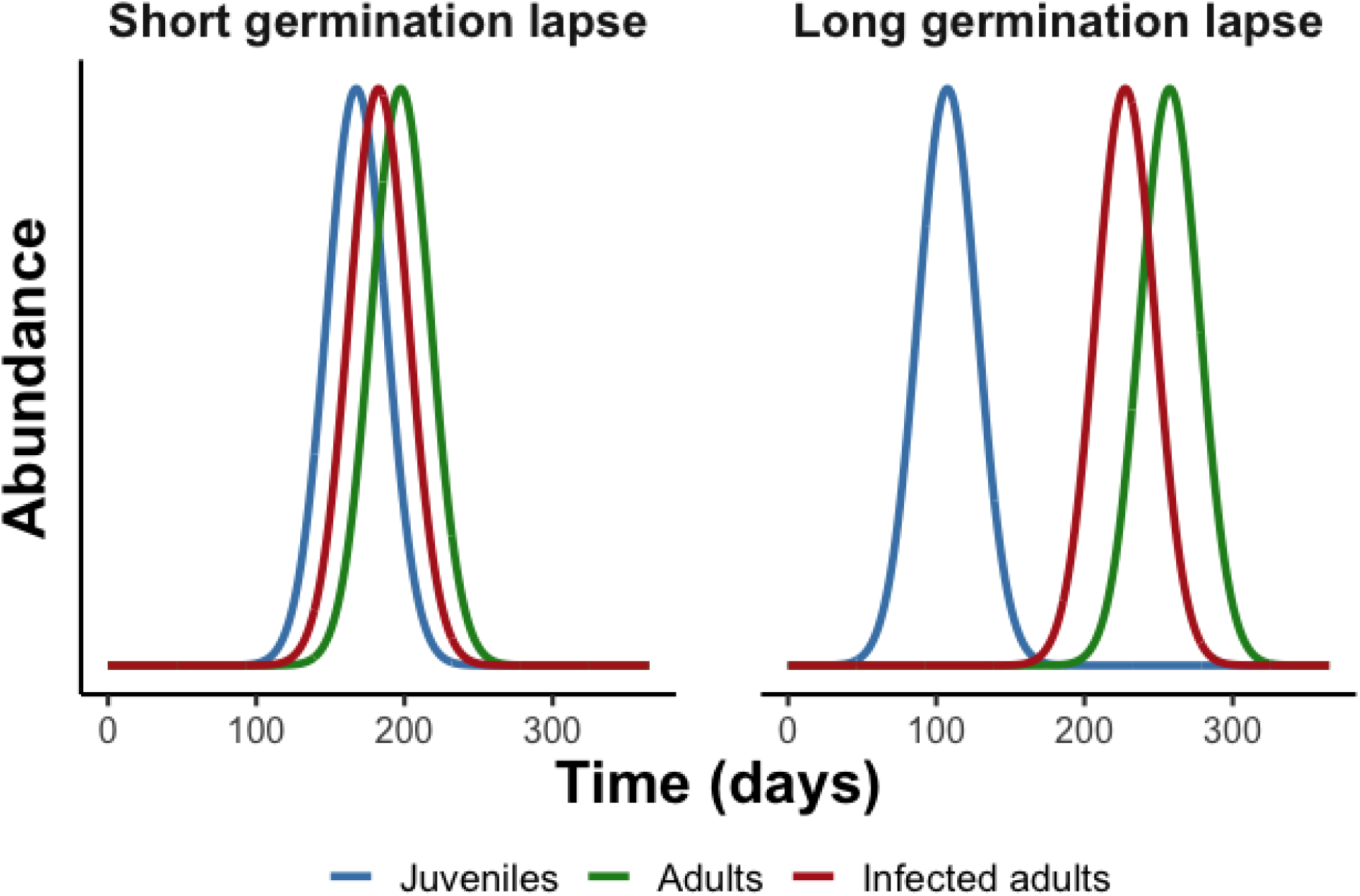
Infectious exploitation of different host age groups. When highly susceptible age groups are further spaced apart due to phenological lapses, the pathogen faces a complex landscape of which age group to exploit.

A clear example of these different transmission opportunities can be illustrated by ‘anther-smut disease’, a pollinator-transmitted fungal pathogen of plants in the Carnation (Caryophyllaceae) family. The fungus produces spores in the anthers of flowers, replacing the pollen and resulting in chronic sterilization of the host. While the fungus resides asymptomatically within meristem tissue within the plant year round, transmission is limited to the flowering season. Transmission is facilitated by pollinators, but can also occur through localized wind dispersal. There is evidence that the pathogen can manipulate the onset of flowering in infected hosts, pushing flowering time either earlier or later than uninfected plants (Bernasconi et al. 2009). In the well-studied species, *Silene latifolia*, infected plants flower a few days earlier than healthy plants, such that infected flowers are present during the peak bloom of healthy plants, and can therefore take advantage of pollinator transmission (M. Hood, unpublished data). However, Tang et al. (2019) found that in several Himalayan species of *Silene*, peak flowering for infected plants was advanced as much as four weeks compared to healthy individuals. At first glance, this is a puzzling transmission strategy, because very few healthy target plants are flowering. One possible explanation is that the early flowering behavior of infected plants is a pathogen manipulation strategy to track peak abundance of the vulnerable juvenile stage, which can be infected through localized wind dispersal. Given the differences in timing for the Himalayan flowers, this raises the question: what aspects of host phenology and physiology can drive selection on the pathogen’s transmission timing?

This question applies to floral pathogens generally. Pathogens that are florally transmitted can transmit only while their hosts are flowering, or are producing flower-like structures. Host flowering phenology therefore determines the pathogen’s transmission window, and several floral pathogens alter its timing. For example, the *Puccinia monoica* rust induces pseudoflowers on *Arabis drummondii* that appear alongside the flowers of another species, *Ranunculus inamoenus*, rather than with healthy *Arabis* (Roy 1993, reviewed in McArt et al. 2014). Floral pathogens therefore offer a natural setting in which to ask how the timing of host life stages shapes the evolution of pathogen transmission timing.

Here, we build a seasonally forced, age-structured, compartmental model of *Silene latifolia* (white campion) and its sterilizing anther-smut fungus, *Microbotryum*, a model floral-pathogen system, to ask how the timing of separate host life stages shapes the prevalence and evolution of pathogen transmission timing. Our work has four goals. First, we quantify how the timing of seed germination affects equilibrium disease prevalence, and how this depends on juvenile susceptibility and maturation rates. We hypothesized that separating germination from flowering would reduce prevalence, providing a mechanism for hosts to avoid pathogen infection. Second, we identify the timing of infectious flowering that maximizes prevalence when germination occurs at different points of the year. We hypothesized that prevalence would be highest when infectious flowering overlaps the most profitable pool of susceptible hosts, which depends on the relative susceptibility of juvenile and adult hosts, contact, and maturation rates. Third, we use invasion analysis to find the evolutionarily stable timing of infectious flowering across germination times and maturation rates. This same approach allows us to test two alternative hypotheses: 1) selection favors a single infectious flowering time that tracks the best opportunity, or 2) temporally separated pools of juvenile and adult hosts maintain a polymorphic population of pathogen genotypes, each exploiting a different host life stage.

Finally, we repeat the evolutionary analysis with parameters for *Dianthus pavonius*, an anther-smut host whose juveniles are much more susceptible relative to adults, and mature much faster than those of *S. latifolia*, to ask whether these life-history differences change the evolutionary outcome.

### The Model

To investigate how host phenology and seasonal dynamics can impact selection on disease transmission timing, we developed an age-structured model based on the *Silene-Microbotryum* system. Our model incorporates key host life-history features such as host age structure and flowering phenology, both of which are seasonally driven. Seasonal variation is represented through time-dependent functions that drive processes such as seed germination, flowering, and disease transmission. Consistent with the biology of anther-smut, we assume infected hosts are permanently sterilized, such that only susceptible hosts contribute to reproduction. We also assume that onward transmission occurs exclusively through spores dispersed from infected flowering individuals. Initially, we assumed uniform, non-evolving host and pathogen populations, however we later incorporated evolutionary invasion analysis in our model.

### Host demographic model

Individuals progress through four developmental stages: seeds (*S*_*d*_), juvenile (*S*_*j*_), vegetative (*S*_*v*_), and flowering (*S*_*f*_) (Figure S1). Seeds germinate into juveniles at a seasonally driven rate *g(t)*, while maturation from juvenile into adult stages occurs at a constant rate *m*. Adults transition between vegetative (*S*_*v*_) and flowering stages (*S*_*f*_) within each season, at seasonally driven rates *v(t)* and *f(t)*. This represents a winter vegetative stage, which moves into a flowering stage as the temperature warms. Plants return to a non-flowering state when the temperature cools.

All compartments experience a constant mortality rate µ, except seeds, which die at a quarter of this rate (μ / 4; Equation 4). Because this mortality is density-independent, population regulation is instead introduced through a density-dependent mortality term proportional to population size (γ*N*), which is only applied to both juvenile groups (*S*_*j*_ and *I*_*j*_) to reflect competition for space and light. This formulation follows established approaches to host–pathogen coexistence in structured populations (Getz & Pickering 1983).

### Seasonal forcing

We modelled temperature as a sine function of time with a period of one year. Seasonal forcing parameters are then functions of temperature, using a specific temperature threshold, κ (25 °C, Table 1), for an on-off switch. The switch is modeled as two opposing logistic curves to remain differentiable. Seasonal forcing occurs through three key processes: germination, *g(t)*, and stage transitions between vegetative and flowering stages, *v(t)* and *f(t)* (Equations 1–3). We develop phase-shifted temperature curves to implement temporal offsets equivalent to the phase offset for any of these functions. These functions reflect temperature-dependent phenological shifts commonly observed in plant populations, including distinct germination and flowering periods. We allow the threshold timings to differ between infected and uninfected individuals, capturing pathogen-induced shifts in flowering phenology. Each seasonal rate peaks for roughly 98 days per year, while its phase-shifted temperature exceeds κ, which is centered on the day of peak temperature for that process. Throughout, we describe the timing of germination, healthy flowering and infectious flowering by the midpoint of this window (the day on which the rate peaks): for example, a flowering midpoint of day 182.5 means plants flower from about day 134 to day 231, with the number of flowering plants peaking near day 182.5.

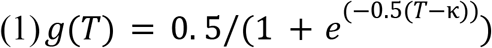

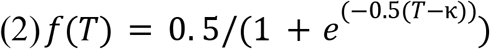

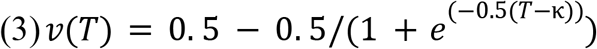

**Table 1.** Model parameters and the values used for *S. latifolia* (Slowinski et al. 2025) and *D. pavonius* (Bruns et al. 2017). *S. latifolia* values are used in all simulations unless stated otherwise.

| Variable | Description | Value (*S. latifolia*, per day) | Value (*D. pavonius*, per day) |
| --- | --- | --- | --- |
| $g(t)$ | Seasonal Germination Rate | Peak of 0.5 | Peak of 0.5 |
| $m$ | Maturation Rate | 0.0023 | 0.022 |
| $v(t)$ | Seasonal Flowering to Vegetative Transition Rate | Peak of 0.5 | Peak of 0.5 |
| $f(t)$ | Seasonal Vegetative to Flowering Transition Rate | Peak of 0.5 | Peak of 0.5 |
| $\beta_j$ | Juvenile Susceptibility Coefficient | 0.0016 | 0.00034 |
| $\beta_v, \beta_f$ | Adult Susceptibility Coefficient | 0.0011 | 0.000034 |
| $\mu$ | Mortality Rate | 0.00099 | 0.00051 |
| $b$ | Birth Rate | 0.022 | 0.0036 |
| $\gamma$ | Density-dependence coefficient | 0.001 | 0.00015 |
| $\kappa$ | Flowering Temperature Threshold | 25 °C | 25 °C |

### Pathogen transmission

Infection by anther smut can occur at all post-seed stages, generating corresponding infected stages (*I*_*j*_, *I*_*v*_, *I*_*f*_). We assume that transmission at all host life stages is driven by contacts with infected flowering individuals (*I*_*f*_). We alter the temperature thresholds to begin flowering between infected and susceptible individuals, which captures a pathogen-induced shift in flowering phenology. This difference generates an effective “infection lapse,” representing the difference in the peak abundances of infectious flowering and healthy flowering plants. These dynamics are described by the following system of non-autonomous ordinary differential equations (Equations 4–10).

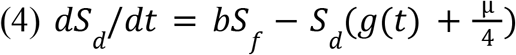

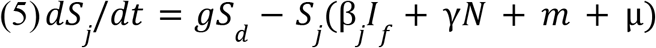

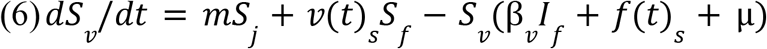

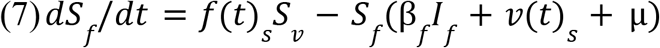

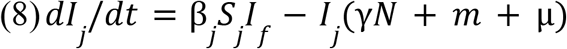

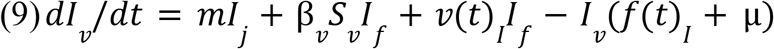

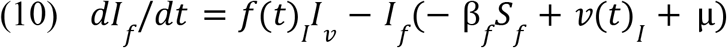

Our model utilizes density-dependent transmission in order to more easily compare transmission at two stages; but for the anther-smut system pollinator transmission to adults is known to be frequency-dependent (Bruns et al. 2017). As is observed in the *Silene-Microbotryum* system, we assume that juveniles are more susceptible to infection than adults (both flowering and vegetative), such that:

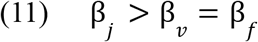

Where β_*j*_, β_*v*_, and β_*f*_ are the transmission coefficients for juvenile, vegetative, and flowering plants respectively. A schematic representation of these dynamics is shown in Figure S1 with a table of parameters in Table 1.

### Simulations

We used our model to investigate four questions about the fitness impacts of seasonal mismatches between hosts and pathogens. First, to measure the impact of temporal overlap between germination and flowering on pathogen prevalence, we investigated the impact of varying host maturation rates post germination on varying germination midpoints. Second, we explored whether varying the infectious flowering timing could lead to a greater disease prevalence. Third, we evaluated evolutionary outcomes using invasion analysis to determine which infectious flowering periods would be favored by selection, and how this depended on host susceptibility and phenology. Finally, we repeated the evolutionary analysis with parameters for *D. pavonius* to ask whether the outcome depends on host life history.

All simulations were run in Python 3.13.7, with the NumPy, SciPy and Matplotlib libraries, using an explicit Runge-Kutta method for solving differential equations. Unless specified, parameters were held constant at the *S. latifolia* values in Table 1 while varying specific host or pathogen parameters across 30 evenly spaced midpoints spanning the year (about every 12.5 days; Figures 2 and 3). The evolutionary analysis (Figures 4–6) used 18 germination midpoints from day 62.5 to day 302.5 (about every 14 days), with infectious flowering midpoints sampled as described below. Model outcomes and pathogen success were determined using equilibrium pathogen prevalence, calculated as the mean proportion of infected individuals over a single year after year 400 (excluding *S*_*d*_, the seed class). We visually assessed the dynamics of the model and determined that 400 years was sufficient for our model to approach a stable equilibrium (Figure S2). For the evolutionary analysis, pathogen success was instead measured by invasion fitness, as described below. We began each simulation with 5 susceptible and 5 infected juvenile individuals.

**Figure 2.**
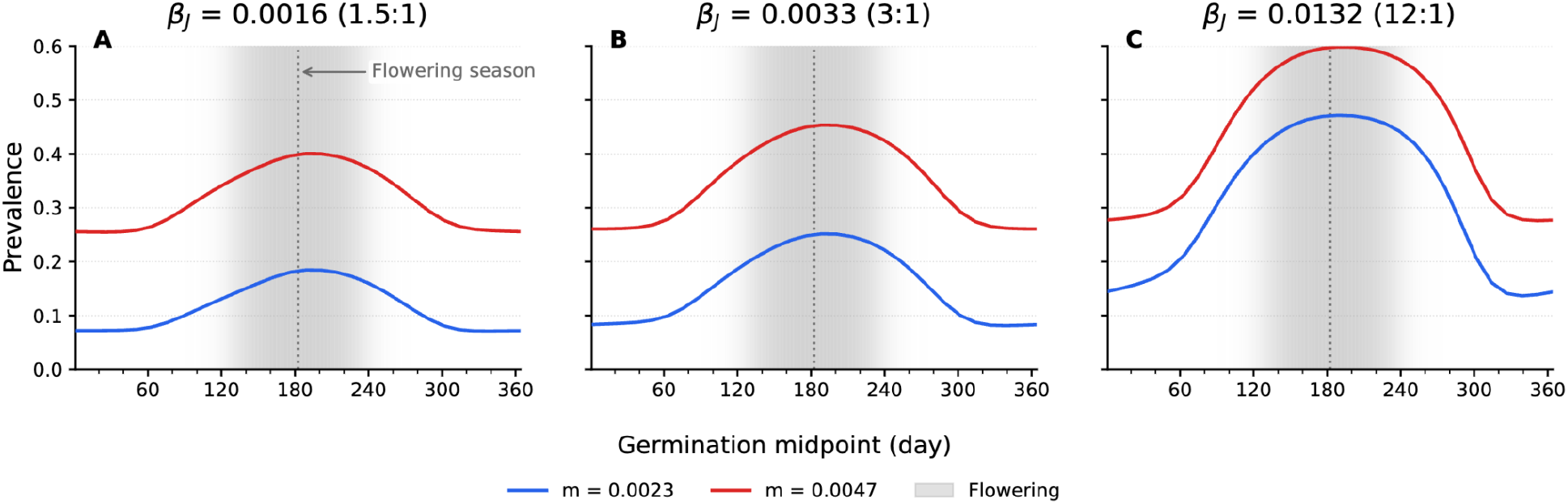
Equilibrium prevalence against germination midpoint for variable juvenile transmission and maturation rates. Disease is highest when germination coincides with flowering, and the effects of germination timing increase with juvenile susceptibility. Gray shading shows healthy flowering (midpoint day 182.5). Other parameters as in Table 1.

**Figure 3.**
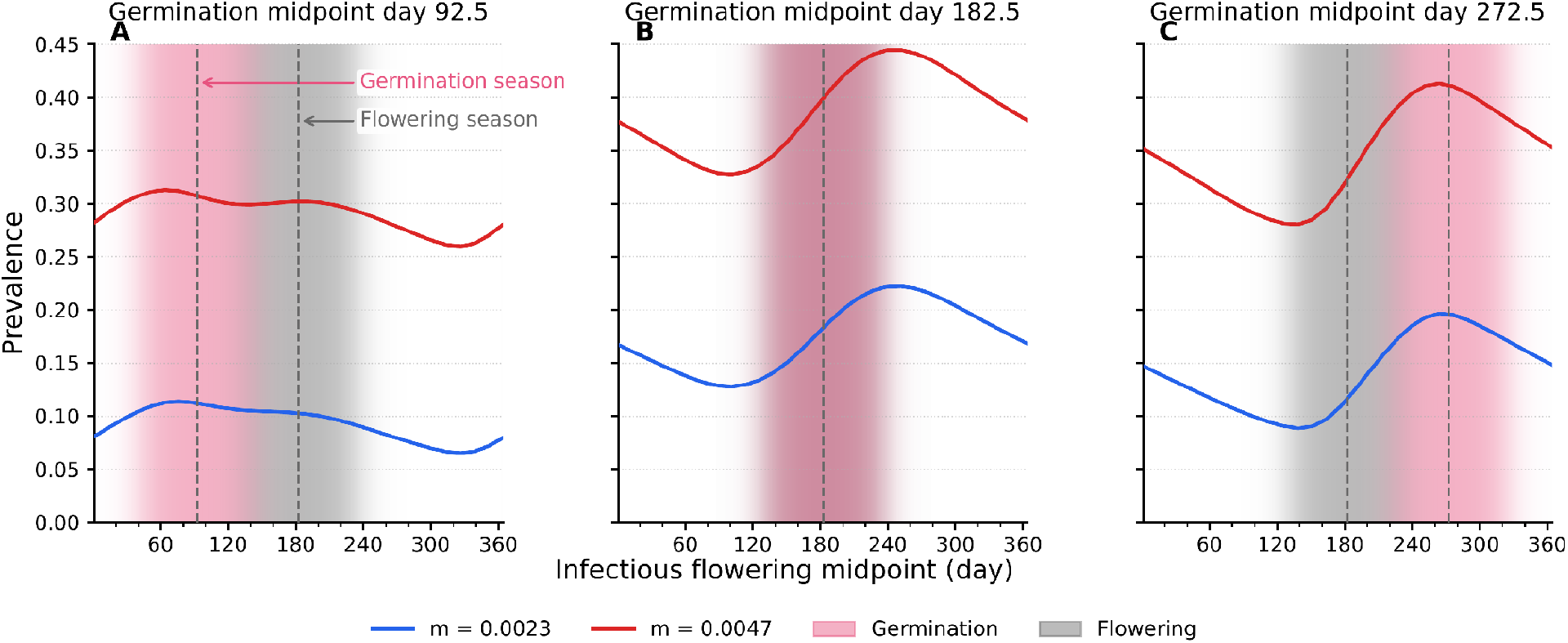
Equilibrium prevalence against infectious flowering for variable germination midpoints and maturation rates. Prevalence peaks when infectious flowering follows the main juvenile cohort. Pink shading shows germination, gray shading shows healthy flowering. Other parameters as in Table 1.

**Figure 4.**
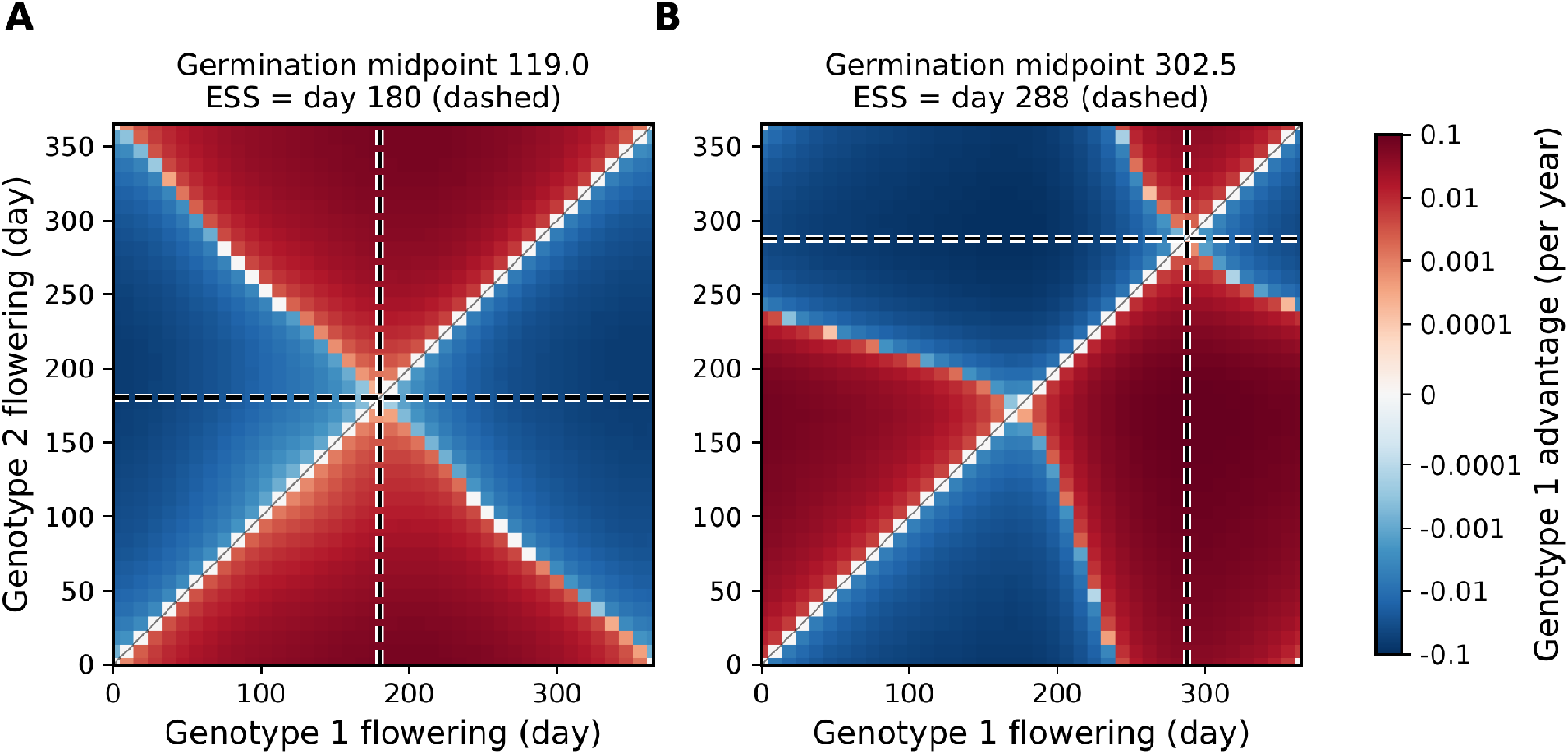
Invasion analysis with two pathogen genotypes differing in infectious flowering midpoint. *S. latifolia* shows a single ESS across germination times. **(A)** Germination midpoint day 119, ESS day 180. **(B)** Germination midpoint day 302.5, ESS day 288. Red shading shows genotype 1 wins, blue shading shows genotype 2 wins, and white shading shows the two coexist (only along the diagonal, where genotypes are nearly identical). The dashed lines show the ESS.

**Figure 5.**
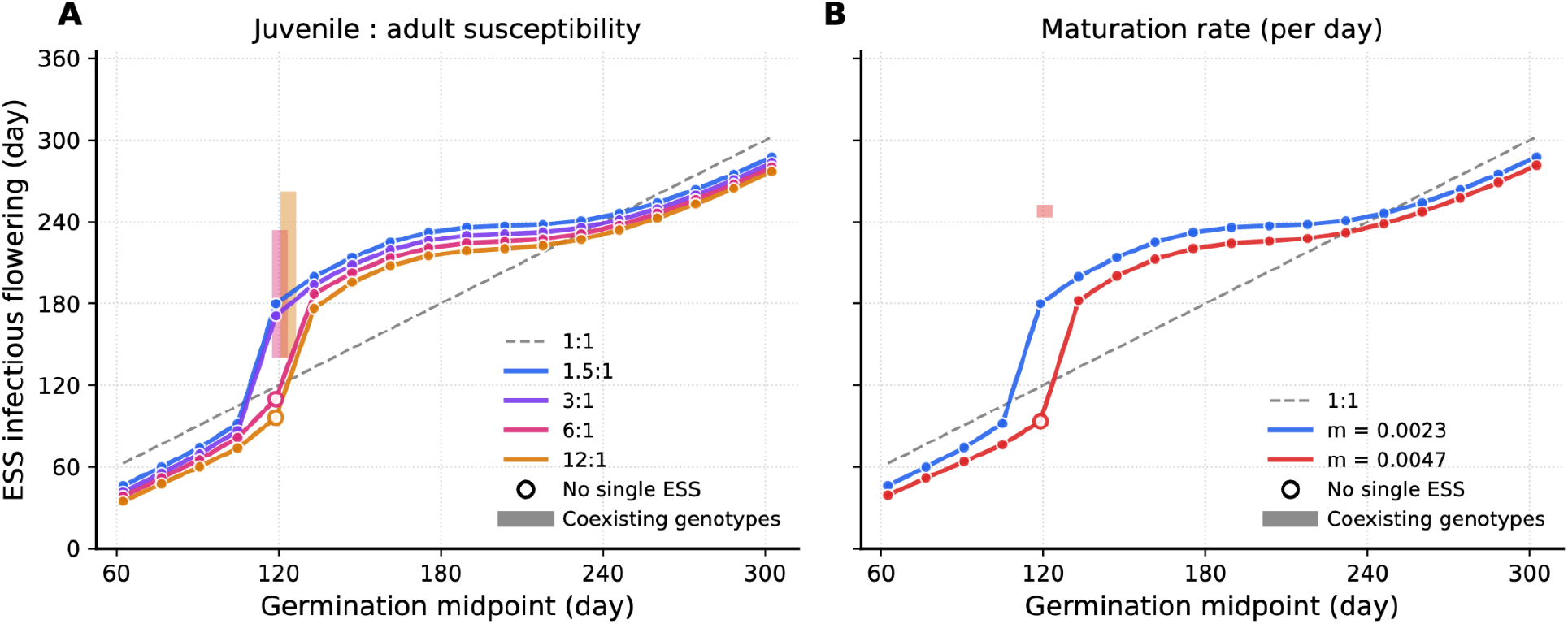
ESS infectious flowering midpoints against germination midpoints. (A) *S. latifolia* parameters with juvenile:adult susceptibility of 1.5:1 (estimated), 3:1, 6:1 and 12:1. (B) *S. latifolia* parameters at the estimated maturation rate (m = 0.0023 per day) and twice that rate. Either high juvenile susceptibility or fast maturation produces polymorphism when germination splits into two cohorts. Filled points show ESS, open circles mark germination midpoints with no single ESS, and the vertical extent of each shaded bar spans the infectious flowering times that form a protected polymorphism with the open-circle strategy. The dashed line shows a 1:1 correlation between germination midpoint and ESS flowering midpoint.

**Figure 6.**
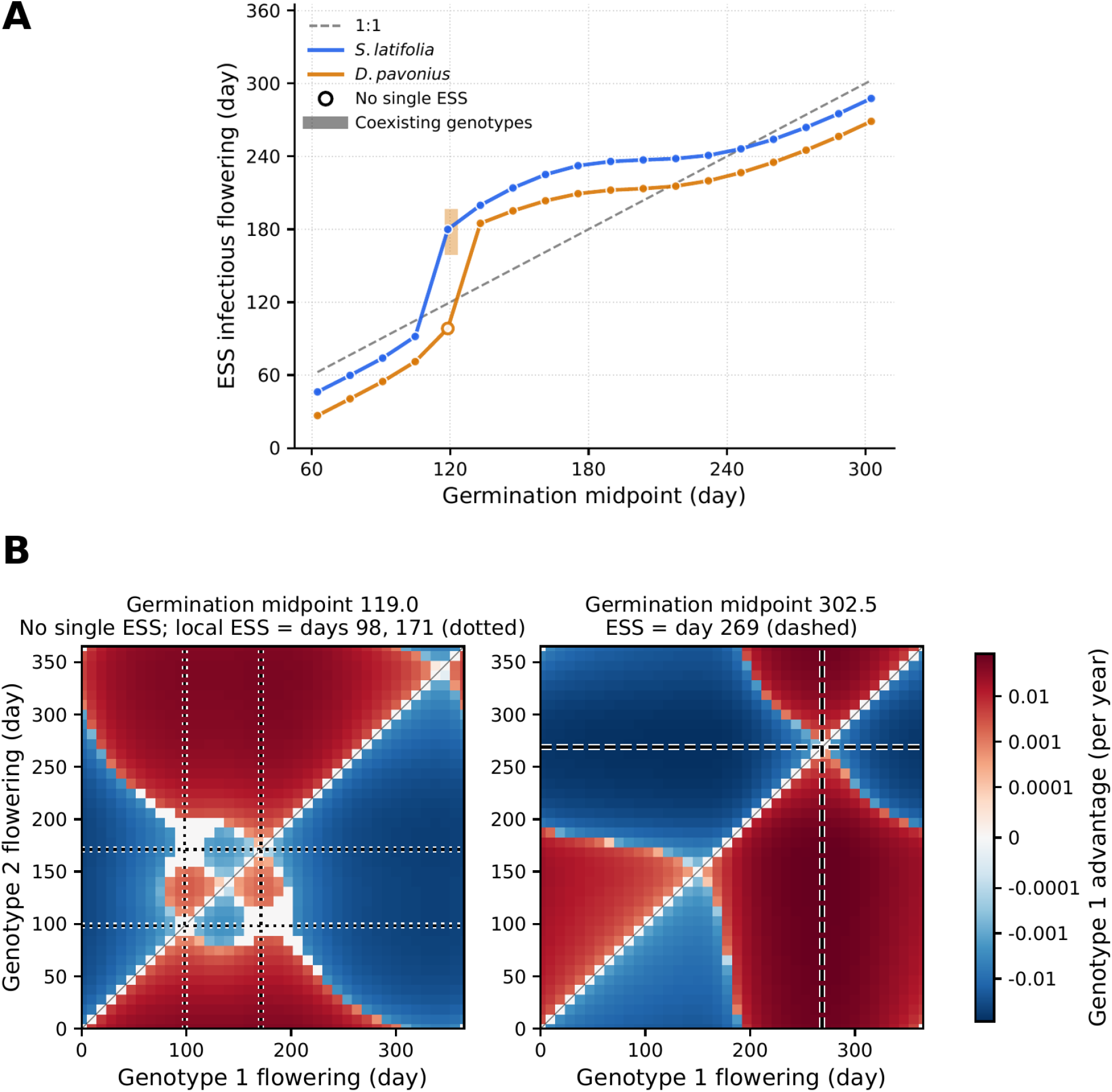
ESS infectious flowering time against germination midpoint between species. Only *D. pavonius* is polymorphic at the estimated parameters. (A) ESS for *S. latifolia* in blue and *D. pavonius* in orange. Open circles mark germination midpoints with no single ESS, the vertical extent of the shaded bar spans the infectious flowering times that can coexist with the open-circle strategy. (B) Competition raster for *D. pavonius*, as in Figure 4. At germination midpoint day 119 no single ESS exists: dotted lines mark the two locally uninvadable strategies (days 98 and 171), and white regions off the diagonal mark pairs of genotypes that each invade the other. The confirmed polymorphism pairs the convergence-stable strategy at day 98.4 with the invading genotype at day 187.2, the flowering time with the strongest mutual invasion. At midpoint 302.5 the dashed lines mark the single ESS (day 269).

### Effect of germination timing on prevalence

One strategy for hosts to reduce pathogen pressure could be to space out vulnerable life history stages for transmission. For example, if the germination is delayed beyond the peak of flowering, there may not be many infectious flowering plants in the population remaining by the time germination occurs. However, this could also limit the growth and reproduction of the host population. To establish a baseline for how germination timing affects infection pressure, we used our model to explore how greater temporal separation between germination and flowering affected the transmission and prevalence of a pollinator-vectored disease. Here we assumed that infected plants were constrained to flower during the same period as uninfected plants. We then varied the midpoint of seasonal germination *g(t)* over the full year (0–365 days), while holding susceptible and infected flowering time at mid-year (day 182.5). All flowering and germination periods were maintained at approximately 98 days long. Additionally, we explored how juvenile susceptibility (β_*j*_) and the maturation rate (*m*) altered the effect of germination timing on infection prevalence. For each parameter combination, we identified equilibrium disease prevalence across the year to determine the influence of birth pulse and flowering timing.

### Effect of infectious flowering timing on prevalence

Next, we examined how temporal separation between host germination and flowering affects selection on pathogen transmission timing. We varied the midpoint of infectious flowering across the year while keeping healthy host flowering constant at day 182.5. Both flowering periods were held constant at approximately 98 days long. We explored how this influence is dependent on certain crucial developmental host life-history parameters such as the timing of peak seed germination, or birth pulse. For each scenario, we determined whether pathogen success was maximized by synchronizing with juvenile birth pulses, adult flowering, or intermediate seasonal windows by treating infection prevalence as a function of the midpoint of infectious flowering.

### Invasion analysis of infectious flowering timing

While single-genotype analyses identify conditions that optimize transmission patterns, they do not capture allelic invasion or competition dynamics. To assess pathogen evolutionary outcomes, we therefore used invasion analysis (Geritz et al. 1998): for each resident genotype with a given infectious flowering time, we asked whether a rare mutant genotype with a different flowering time could invade. An evolutionarily stable strategy (ESS) is a resident flowering time that no mutant can invade, and a protected polymorphism is a pair of flowering times that can each invade the other. This analysis allowed us to distinguish between two potential pathogen evolutionary trajectories: convergence on a single transmission timing, or maintenance of a polymorphism of different transmission timings.

For each pair of genotypes, we first ran the model with the resident genotype alone until equilibrium. We then asked whether a second, mutant genotype introduced at low frequency could invade. While rare, the mutant has no effect on the resident, so its infected classes follow a linear system whose coefficients repeat every year. The long-run per-year growth rate of this system, or the mutant’s invasion fitness (*s*(*resident, mutant*)), shows invasion when *s* > 0 (Klausmeier 2008). We computed *s* for all pairs of 40 infectious flowering midpoints spanning the year, identified singular strategies where the selection gradient is zero, refined each strategy with root-finding, and tested it against mutants at one-day resolution across the whole year.

Convergence-stable strategies that no mutant could invade were classified as ESS for the respective germination time. Pairs of genotypes that could each invade the other (*s* > 0 in both directions) were classified as protected polymorphisms. Because invasion fitness describes only a rare invader, we confirmed each candidate polymorphism by simulating the full two-genotype model. We paired the convergence-stable flowering time with the invading flowering time, ran each genotype alone to its annual cycle, introduced the other at a density of 0.001 infected flowering plants, and simulated for 18,000 years. A polymorphism was confirmed when both runs kept both genotypes and converged to the same share of infected hosts (within 0.001). This approach allowed us to identify ESS and evaluate the potential for polymorphism in the onset of infectious flowering.

Unless stated otherwise, all analyses used parameter values for *S. latifolia* (Table 1). To separate the roles of juvenile susceptibility and maturation, we repeated analyses under *S. latifolia* parameters with the juvenile transmission coefficient raised two-, four- and eightfold (juvenile susceptibility ratio of 3:1, 6:1 and 12:1, with adult susceptibility unchanged) and, separately, with the maturation rate doubled. To test whether the evolutionary outcome depends on host life history, we repeated the invasion analysis with parameter values for a second anther-smut host, *Dianthus pavonius* (Bruns et al. 2017; Table 1). *Silene latifolia* is a short-lived perennial, with a modest difference between juvenile and adult susceptibility (Slowinski et al. 2025, Bruns et al. 2022). In contrast, *Dianthus pavonius* is a long-lived perennial, with much greater difference in juvenile and adult susceptibility.

## Results

### Effect of germination timing on prevalence

We first evaluated how the timing of host germination pulses influences pathogen prevalence. Overall, pathogen prevalence was consistently dependent on the timing of the germination pulse, with the maximum prevalence occurring when germination coincided with healthy flowering.

Across all values of juvenile susceptibility and maturation rate, equilibrium prevalence peaked at a germination time of day 189 in all six simulation conditions within one sampling interval (12.5 days) of peak flowering at day 182.5 (Figure 2). Since germination and flowering overlap in these experiments, new juveniles appear while infected plants are flowering. As a result of the slow maturation rate of *S. latifolia*, new juveniles remain susceptible to infection for the rest of the flowering season.

As we increased juvenile susceptibility, we saw an increased average infectious prevalence across the year. This trend is most noticeable when germination overlapped flowering, and under slow maturation (compare Figure 2A and 2C). Separating germination from flowering reduced prevalence by 36–71%, with the largest reductions under slow maturation. Counterintuitively, faster maturation increased prevalence. Because infected juveniles cannot transmit until they mature and flower, faster maturation shortens the delay between juvenile infection and onward transmission.

### Effect of infectious flowering timing on prevalence

We next tested how variation in the timing of infectious flowering affects pathogen prevalence. Pathogen prevalence depended heavily on host life-history dynamics, with peak prevalence occurring when infectious flowering coincided with high juvenile abundance. The timing of peak equilibrium prevalence shifted depending on the relative timing of the germination pulse and healthy flowering, rather than occurring at a fixed point in the infectious flowering cycle (Figure 3). For germination midpoints of day 92.5, 182.5 and 272.5, juvenile abundance peaked on about day 44, 229 and 232, and prevalence was highest when infectious flowering was centered three to five weeks later: on days 64–76 (earlier for faster maturation), day 251, and day 264, respectively. This reflects the timing of susceptible juveniles relative to susceptible adults.

However, this pattern was also influenced by the alignment between germination and healthy flowering timing and the maturation rate. Specifically, an increased maturation rate reduced the relative impact of infectious flowering timing on infection prevalence, as this led to a shorter juvenile stage, and therefore less opportunity to infect juveniles. This can be seen in Figure 3B: infectious prevalence is suppressed when infectious flowering precedes germination. Variation in germination timing also manipulated the location of the prevalence peaks, reflecting interactions between the birth pulses and both infected and susceptible flowering times. Despite these shifts, pathogen success was always highest when infectious flowering overlapped the period of greatest juvenile abundance, during or shortly after the germination pulse.

### Evolutionarily stable infectious flowering timing

To determine if pathogens with different transmission timings could co-occur in a stable polymorphism, we used invasion analysis with two distinct pathogen genotypes differing in their flowering time. Given the *S. latifolia* parameters in Table 1, a single infectious flowering time was evolutionarily stable at every germination midpoint we examined (18 midpoints from day 62.5 to day 302.5). For example, for a germination midpoint of day 119 the ESS was day 179.9, close to the peak of healthy flowering (day 182.5), and for a midpoint of day 302.5 it was day 287.6 (Figure 4). Measured against median seed germination time, the ESS was highly consistent: at every germination midpoint, the evolutionarily stable infectious flowering period was centered 44–60 days after the median germination day.

We next examined how this ESS flowering timing depended on germination timing and the host maturation rate. We found a positive nonlinear association with the germination timing and the ESS pathogen flowering timing (Figure 5), with variation across maturation rates. At the *S. latifolia* maturation rate, the ESS preceded the germination midpoint when the midpoint was early (ESS at day 46 for a germination midpoint of day 62.5) or late (ESS day 288 for a midpoint of day 302.5), but switched abruptly to near or after the peak of healthy flowering for midpoints between about day 119 and day 240 (ESS day 236 for a midpoint of day 189.6). The switch occurred where germination changes from a spring flush of over-wintered seeds to a summer cohort germinating as seeds are produced (Figure S3). The ESS remained 44–60 days after the median germination day at every germination midpoint.

When tested with twice the *S. latifolia* maturation rate, the ESS shifted 6–18 days earlier and delayed the switch to later germination midpoints. In these conditions, no single flowering time was evolutionarily stable at a germination midpoint of day 119, where germination is split almost equally between the spring and summer cohorts. Simulating the full two-genotype model confirmed a polymorphic population: genotypes flowering at days 93.4 and 252.7 converged to the same equilibrium, with 57% and 43% of infected hosts respectively, from either starting condition (18,000 simulated years; open circle and bar in Figure 5B). This indicates that evolutionary outcomes in pathogen transmission timing are strongly shaped by host seasonal dynamics and life history traits such as maturation. Together, these results demonstrate that pathogen transmission timing evolves in response to host phenology, with a dominant strategy emerging under most conditions, and a protected polymorphism in a narrow window of germination timing under specific conditions.

We also tested the model by raising juvenile susceptibility (Figure 5A), which shifted the ESS earlier at every germination midpoint, toward the timing of juvenile availability (by 4–9 days at a juvenile:adult ratio of 3:1 and 11–18 days at 12:1), without changing the shape of the relationship. At 6:1 and 12:1 no single flowering time was evolutionarily stable at a germination midpoint of day 119, and simulations of the full two-genotype model confirmed a protected polymorphism (genotypes flowering at days 109.8 and 215.3 at 6:1, and 96.2 and 234.0 at 12:1) coexisting from either starting condition.

Lastly, we compared the change in ESS over germination time between our *S. latifolia* parameterization and *Dianthus pavonius* parameterization, an anther-smut host whose juveniles are ten times more susceptible than adults (compared with 1.5 times for *S. latifolia*) and mature about ten times faster (Table 1). We found that the systems modeled around *Silene* and *Dianthus* retained a strong relationship between ESS flowering time and germination time (Figure 6A). However, under *D. pavonius* parameters, no single flowering time was evolutionarily stable at a germination midpoint of day 119 (Figure 6B), where germination is split almost equally between a spring flush of over-wintered seeds (53% before day 130) and a summer cohort germinating as seeds are produced. Simulating the full two-genotype model at this germination time confirmed a protected polymorphism at days 98.4 and 187.2, with 36% and 64% of infected hosts respectively. One genotype therefore transmits to spring-germinated juveniles, while the other transmits among flowering adults and summer-germinated juveniles. At every other germination midpoint *D. pavonius* had a single ESS (e.g., day 268.8 for a midpoint of day 302.5; Figure 6B), centered 23–36 days after the median germination day, earlier than in *S. latifolia* (44–60 days), consistent with its faster maturation. These results show that the dependence of the pathogen’s optimal transmission timing on host life history holds under a second, independently estimated parameter set.

## Discussion

Our results highlight the importance of seasonal variation in host age structure in determining how pathogens optimally transmit. We found that pathogen prevalence was constrained by the timing of birth pulses and host flowering. In particular, shifting germination away from flowering reduced disease prevalence by up to 71. However, extreme separation between birthing pulses and mating is not always possible for hosts. We varied germination timing across the full year for completeness, but not all of these timings are biologically realistic. When seasonal constraints limit when seedlings can germinate, hosts may have little scope to escape disease by shifting germination. Our results show that even when hosts can fully shift germination across the entire year, pathogen evolution is generally expected to favor a single infectious period, except when germination itself occurs in separate pulses.

Rather than maximizing transmission at a fixed point in time, pathogens must balance exploiting highly susceptible juvenile hosts and highly abundant reproductive adults to find optimal strategies for prevalence. Despite these two discrete pools of susceptible hosts, we found that selection consistently favored a single pathogen genotype at the estimated *S. latifolia* parameters whose infectious flowering largely tracked the timing of juvenile availability, especially when germination and healthy flowering were separated. However, under *D. pavonius* parameters or in *S. latifolia* with faster maturation or higher juvenile susceptibility (see below), polymorphism was found. Across our simulations (Figures 2–6), the infectious flowering time that maximized prevalence, and the ESS, depended on when germination occurred relative to healthy flowering. Generally, this finding demonstrates that optimal transmission timing is shaped by developmental stages, life-history traits, and host availability in a complex landscape of host-pathogen interactions.

The pathogen-centric analyses further show how transmission strategies depend on host life history. Prevalence was highest when infectious flowering was centered three to five weeks after the peak of juvenile abundance (Figure 3), and the ESS lay 44–60 days after the median germination day at every germination time we examined (Figure 5): the pathogen tracks the arrival of susceptible juveniles with a lag, rather than synchronizing with the birth pulse or with healthy flowering. Faster maturation weakened the dependence of prevalence on infectious flowering timing, because a shorter juvenile stage leaves less opportunity to infect juveniles. This lagged tracking offers an explanation for the Himalayan *Silene* populations in which infected plants flower up to four weeks before healthy plants (Tang et al. 2019). Where seedlings emerge well before the healthy bloom, selection favors infectious flowering that arrives early with them. Similar shifts in the timing of infectious structures occur in other floral pathogens.

For example, *Arabis drummondii* plants infected with the rust *Puccinia monoica* produce pseudoflowers at the same time as *Ranunculus inamoenus*, which they resemble, rather than flowering with healthy hosts (Roy 1993, reviewed in McArt et al. 2014). Our framework may therefore apply more broadly to pathogens whose transmission depends on host flowering. For such pathogens, our results predict that the timing of infectious structures should track the arrival of the most susceptible hosts rather than the peak of healthy flowering, and that pathogen populations with more than one flowering time are most likely where juveniles are much more susceptible than adults or mature quickly, and where host germination is split between distinct cohorts.

Pathogens that specialize on either adults or juvenile hosts are commonly observed in nature. Specialization, however, can arise through a single ESS or two coexisting or polymorphic genotypes. One example that could lead to this adaptation would be temporal separation between host age classes. If phenology alone could lead to genetic polymorphisms, particular pathogen genotypes would then be under selection to further maximize their age-specific transmission, taking advantage of the susceptibility of different host age classes. Alternatively, if this separation in time does not promote polymorphism, it may still provide an avenue for a single ESS. We found that when juveniles matured faster or were much more susceptible than adults (in *D. pavonius* and select conditions for *S. latifolia*), temporally separate pools of susceptible hosts maintained coexisting pathogen genotypes, or polymorphism, within a narrow range of host phenologies. However, pairwise invasion analyses revealed that the system contained a single ESS for infectious flowering time in most conditions. Polymorphism arose where germination was split equally between a spring flush of over-wintered seeds and a summer cohort germinating as seeds were produced (Figure 6A). At this germination time, under specific host conditions, we found two groups of pathogens that exploited temporally separate pools of hosts. At the estimated *S. latifolia* maturation rate, the same split in germination produced an abrupt switch of the ESS from the spring to the summer cohort. In *S. latifolia*, polymorphism arose either when maturation was doubled or when juveniles were at least six times as susceptible as adults, but not at its estimated values (slow maturation, 1.5:1 susceptibility). Fast maturation keeps each juvenile cohort brief, and high juvenile susceptibility makes each cohort valuable, which strengthens the effect of the spring and summer cohorts as distinct transmission opportunities worth specializing on. Away from this window, one transmission opportunity dominates and a single genotype is favored.

Two features may limit how readily this polymorphism arises in nature: first, it reflects ecological coexistence rather than evolutionary branching, so gradual evolution through small changes in flowering time cannot split a single pathogen population into two. The second genotype must instead differ in infectious flowering time by roughly 90–160 days, arising realistically through standing variation, immigration, or a mutation with a large effect on phenology. Second, selection within this window is weak, and the polymorphism took thousands of years to establish in our simulations. Importantly, we found that the location of the ESS was strongly dependent on host germination timing, demonstrating that pathogen evolutionary strategies are strongly tied to host phenology as well.

Several extensions of this model and study system could further improve its biological realism and improve its generalizability but increase its complexity and computational cost as well. First, we assume a single transmission pathway across all host life stages, which may be more complex in other biological systems. For example, pathogens that primarily infect juveniles often exhibit density-dependent transmission modes, whereas those that infect adults often exhibit frequency-dependent transmission modes. Incorporating age-specific transmission modes could introduce additional complexity and tradeoffs into the landscape of optimal timing strategies. Secondly, our invasion analysis considers a single trait and two genotypes at a time.

Once a polymorphism is established, both genotypes continue to evolve, and a two-genotype (coalition) analysis would determine where the coexisting pair ultimately settles and whether additional genotypes could invade. Third, our model does not include a latent period between infection and spore production. Susceptible flowering plants that become infected begin transmitting immediately, whereas infected juveniles cannot transmit until they mature and flower. This asymmetry may inflate the benefit of infecting flowering adults relative to juveniles. Because anther-smut infections acquired during flowering may not produce spores until later in the season or the following year, adding a latent period would likely reduce the payoff of adult infection and could shift the ESS further toward the timing of juvenile availability. Fourth, germination in our model responds gradually to temperature, so over-wintered seeds germinate well before the midpoint of the germination window, and the realized timing of germination depends on when seeds are produced. Explicit germination cues or seed dormancy would give hosts more precise control over germination timing, and could change how often germination is split between spring and summer cohorts, the condition under which we found pathogen polymorphism. Additionally, if the model was structured to allow pathogens to maximize infectivity to either juvenile or adult hosts, polymorphism may be more likely.

Together, our results offer a framework for understanding how seasonal structure shapes pathogen evolution. We show that pathogen transmission timing shows a quantifiable yet complex relationship dependent on maturation rates, susceptibility ratios, and other host life-history parameters. In systems such as *Silene-Microbotryum*, shifts in host phenology can have significant effects on pathogen fitness and evolutionary trajectories. More broadly, our findings suggest that predictable seasonal processes can act as strong selective forces on pathogen strategies. Where juveniles mature quickly or are much more susceptible than adults, they can also maintain more than one pathogen strategy at once.

## Acknowledgements

We thank the members of the Bruns Lab at the University of Maryland for their feedback, and Professor Michael Hood at Amherst College for sharing unpublished observations on flowering phenology in *Silene latifolia*.

## Code availability

All model code and the scripts that reproduce every figure and analysis are available at https://github.com/bds062/Host-Pathogen-Evolutionary-Dynamics-Model.

## Ethics statement

Our study involved no human participants, animals, field collections or regulated materials.

