## Supplemental Document for "When should a pathogen strike? Navigating the landscape of birth pulses and mating season"

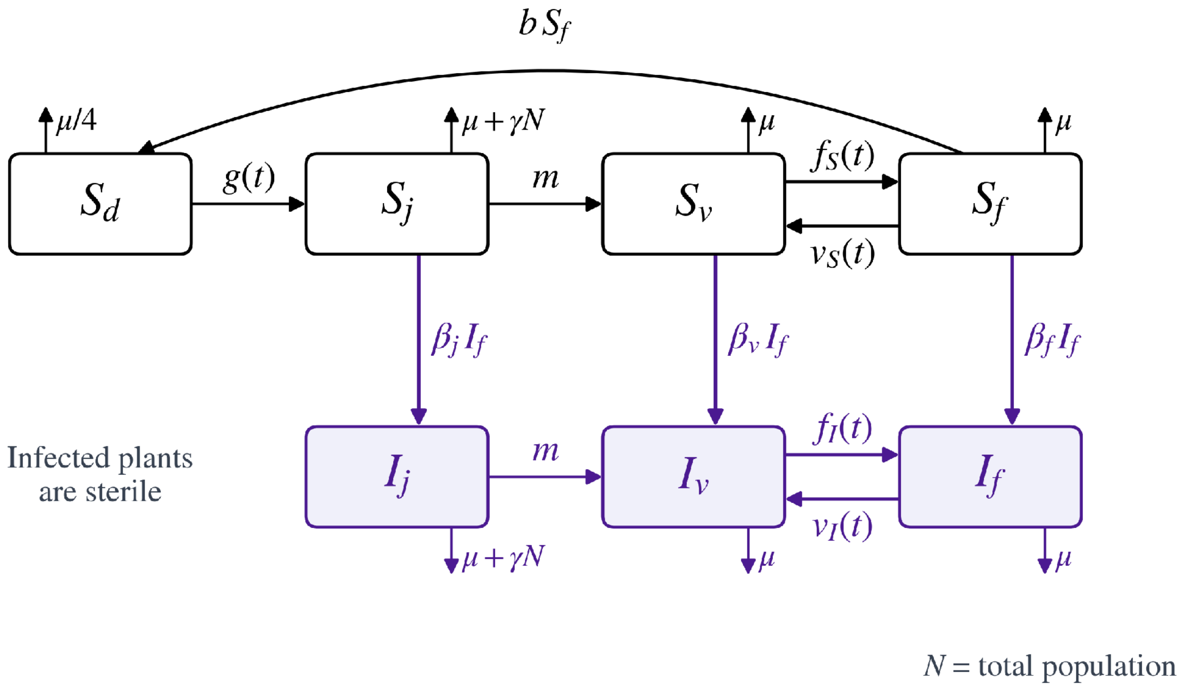


**Figure S1. Structure of the age-structured host–pathogen model.** Susceptible plants pass from seeds ($S_d$) to juveniles ($S_j$), vegetative adults ($S_v$) and flowering adults ($S_f$). Germination $g(t)$ and reproductive stage transitions, $f(t)$ and $v(t)$, are seasonally forced. Infected plants ($I$) use their own flowering thresholds. Juveniles mature at a constant rate $m$. Infection (purple) occurs by contact with infectious flowering plants $If$ at stage-specific rates $\beta_j$, $\beta_v$, and $\beta_f$. Only healthy flowering plants produce seeds (rate $b$). Infected plants are sterile. All stages die at rate $\mu$ (seeds at $\mu/4$), and juveniles additionally experience density-dependent mortality $\gamma N$. Parameter values are given in Table 1 of the main text.

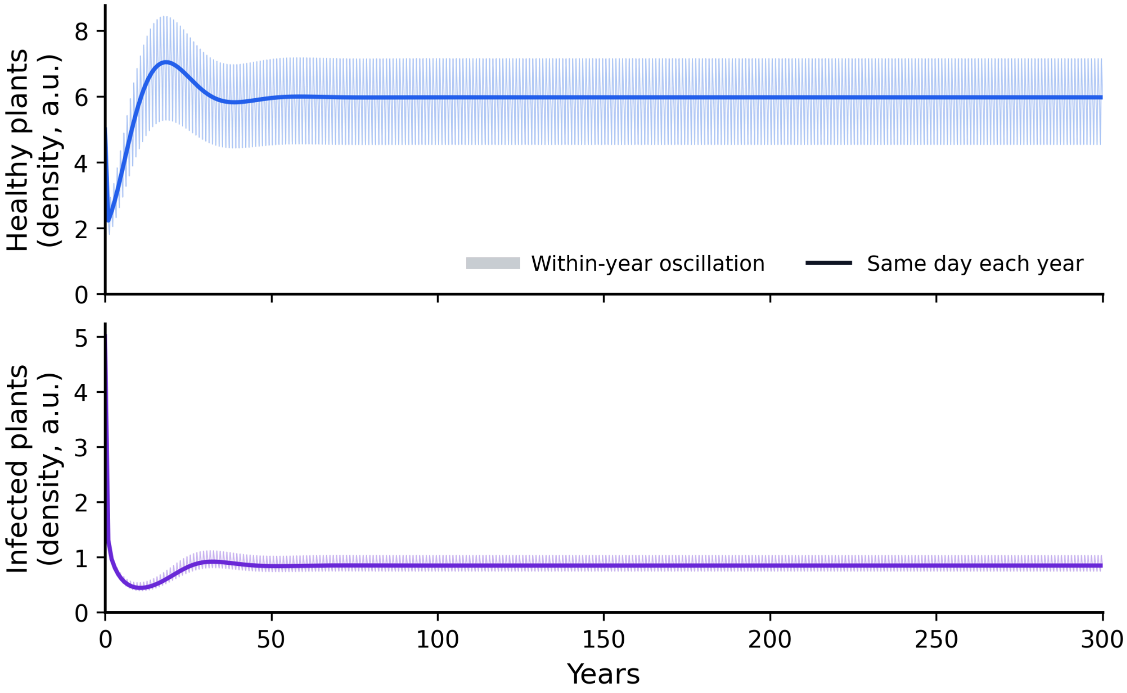


**Figure S2. The model reaches a repeating annual cycle well within 400 years.** Total density of healthy (top) and infected (bottom) plants, in the arbitrary units of the model, over 300 years from the initial condition used in all simulations (5 susceptible and 5 infected juveniles), under *S. latifolia* parameters with germination midpoint day 119 and infectious flowering at its ESS (day 180).

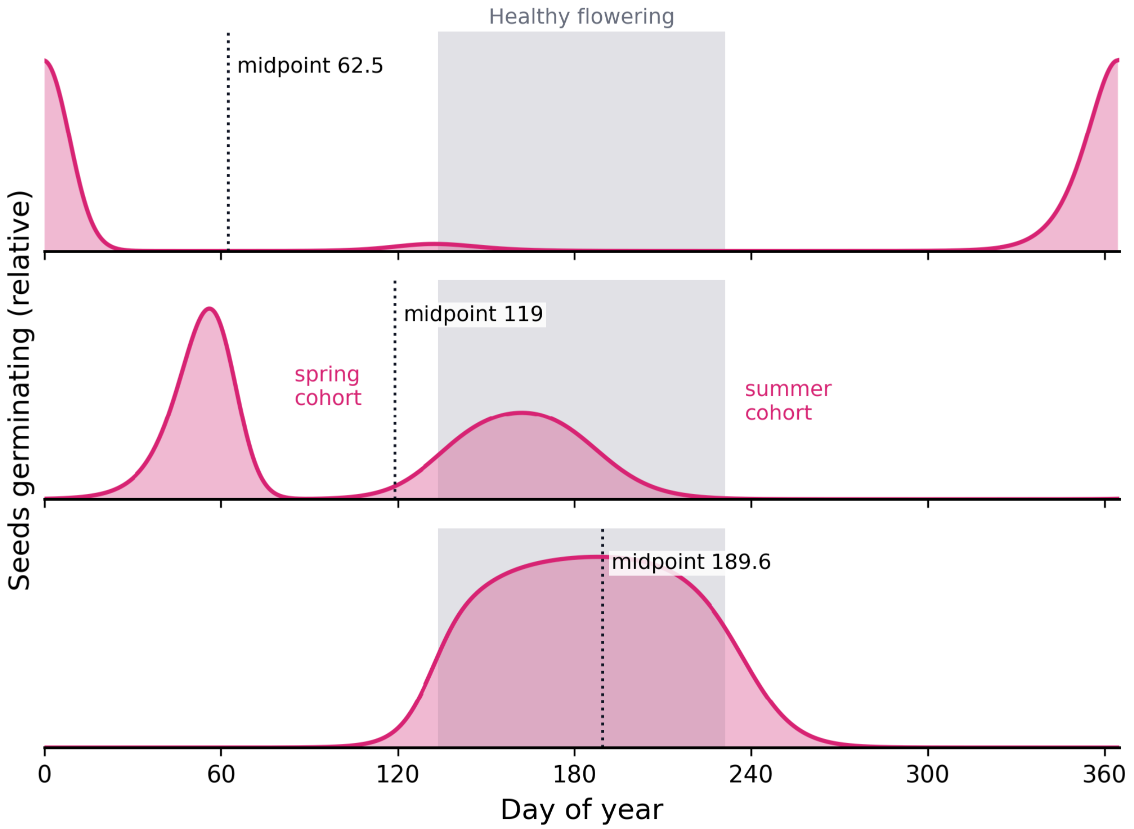


**Figure S3. Realized germination timing for three germination midpoints.** Daily germination under *S. latifolia* parameters, with infectious flowering at the ESS for each midpoint. For an early midpoint (day 62.5) almost all germination occurs around beginning/end of the year. For an intermediate midpoint (day 189.6) seeds germinate during flowering. For a midpoint between the two, (day 119) germination is split almost equally between a spring and summer cohort. This is the germination timing at which polymorphism was found. Gray shading shows healthy flowering (midpoint day 182.5).
